# Human engineered heart tissue transplantation in a guinea pig chronic injury model

**DOI:** 10.1101/2021.06.28.450176

**Authors:** Constantin von Bibra, Aya Shibamiya, Birgit Geertz, Eva Querdel, Maria Köhne, Tim Stuedemann, Jutta Starbatty, Arne Hansen, Bernhard Hiebl, Thomas Eschenhagen, Florian Weinberger

**Author notes:** equal contribution (Constantin von Bibra, Aya Shibamiya). **Corresponding Author**: Florian Weinberger, MD, Department of Experimental Pharmacology and Toxicology, University Medical Center Hamburg-Eppendorf, Germany. Martinistr. 52, 20246 Hamburg.

## Abstract

Myocardial injury leads to an irreversible loss of cardiomyocytes (CM). The implantation of human engineered heart tissue (EHT) has become a promising regenerative approach. Previous studies exhibited beneficial, dose-dependent effects of human induced pluripotent stem cell (hiPSC)–derived EHT patch transplantation in a guinea pig model in the subacute phase of myocardial injury. Yet, advanced heart failure often results from a chronic remodeling process. Therefore, from a clinical standpoint it is worthwhile to explore the ability to repair the chronically injured heart. In this study human EHT patches were generated from hiPSC–derived CMs (15×10^6^ cells) and implanted epicardially four weeks after injury in a guinea-pig cryo-injury model. Cardiac function was evaluated by echocardiography after a follow-up period of four weeks. Hearts revealed large transmural myocardial injuries amounting to 27% of the left ventricle. EHT recipient hearts demonstrated compact muscle islands of human origin in the scar region, as indicated by a positive staining for human Ku80 and dystrophin, remuscularizing 5% of the scar area. Echocardiographic analysis demonstrated no significant difference between animals that received EHT patches and animals in the control group (fractional area change 36% vs. 34%). Thus, EHT patches engrafted in the chronically injured heart but in contrast to the subacute model, grafts were smaller and EHT patch transplantation did not improve left ventricular function, highlighting the difficulties for a regenerative approach.

## Introduction

Ischemic heart disease is a major cause of death worldwide^1^. Myocardial injury results in an irreversible loss of cardiomyocytes followed by scar formation and remodeling of the remote myocardium, including interstitial fibrosis and cardiomyocyte hypertrophy^2^. There is no curative therapeutic option besides heart transplantation for advanced heart failure patients. Yet, donor shortage restricts its use to a small number of patients and this approach requires lifelong immunosuppression^3^. New treatment options are therefore urgently needed. Human pluripotent stem cell-derived CM (hiPSC-CM) transplantation has become an auspicious approach for cardiac regeneration^4–7^. Transplantation of in vitro grown cardiac tissues exhibited beneficial effects in small animal models^4,8^. Late preclinical and first clinical trials with this approach are ongoing. However, whereas most preclinical trials focused on subacute models^4,5,8,9^, terminal heart failure mostly results from a chronic remodeling process. From a clinical point of view, it is therefore desirable to evaluate the EHT transplantation in the chronically injured heart. Preclinical studies indicate that the grafting of hiPSC-CM injections are less effective in chronically injured hearts^10^. We therefore investigated EHT patch transplantation in a chronic myocardial injury guinea pig model.

## Materials and Methods

### Differentiation of hiPSC-CM and generation of EHT

hiPSCs (UKEi1-A) were differentiated into CMs as previously reported^11^. UKEi1-A was reprogrammed with a Sendai Virus strategy (CytoTune iPS Sendai Reprogramming Kit, ThermoFisher). These studies were approved by the Ethical Committee of the University Medical Center Hamburg-Eppendorf (Az. PV4798/28.10.2014 and Az. 532/116/9.7.1991). EHT patches (1.5×2.5 cm) were cast from cells and a fibrinogen/thrombin mix as recently described^4^ containing 15×10^6^ hiPSC-CMs in 1.5 ml. Control animals received patches generated from the fibrinogen/thrombin mix without cells.

### Study design

24 animals were included in the study. 3 animals died before EHT patch transplantation and 3 animals died during or shortly after EHT patch transplantation. Mortality over the complete duration of the study was 25%. Two animals which showed no decrease in left ventricular function four weeks after injury were excluded from further analysis. All other animals were included. Animals were randomly assigned to the treatment groups and marked with a non-identifying number by an independent investigator. Group size was determined by power calculations based on our previous studies.

### Echocardiography

To evaluate cardiac function transthoracic echocardiography was performed with a Vevo 3100 System (VisualSonics) as previously described^4^. Echocardiographic scans were performed at baseline, 4 weeks after cryoinjury and 4 weeks after transplantation.

### Guinea pig chronic injury model

Female Dunkin Hartley guinea pigs (Envigo, Netherlands) weighing 350-600 g were used as experimental animals. All procedures were performed in accordance with the Guide for the Care and Use of Laboratory Animals published by the National Institutes of Health (Publication No. 85-23, revised 1985) and were approved by the local authorities (Behörde für Gesundheit und Verbraucherschutz, Freie und Hansestadt Hamburg: 109/16). Guinea pigs received buprenorphine (0.05 mg/kg) and carprofen (5 mg/kg) subcutaneously 30 min prior to surgery. Additionally, atropine (0.05 mg/kg) was administered before the induction of anesthesia with isoflurane. Tracheotomy was performed and the guinea pigs were mechanically ventilated (5% isoflurane). Left lateral thoracotomy was executed using the 5th intercostal space. The pericardium was partially removed. For the induction of the cryoinjury a liquid nitrogen cooled probe (5 mm diameter) was applied 3 times for 30 s each to the free left ventricular wall. During surgery the animals received a subcutaneous bolus of saline/electrolyte solution (20 ml/kg). Analgesics (buprenorphine and carprofen) were given for 3 postoperative days. Four weeks after the induction of the cryoinjury the animals underwent a re-thoracotomy and the exposed cryo-lesion was covered with an EHT-patch (15×10^6^ hiPSC-CM), sutured onto the healthy myocardium. Control animals received a cell-free patch. For this, animals were assigned in two experimental groups. Guinea pigs were immunosuppressed beginning 2 days prior to transplantation subcutaneously with cyclosporine (7.5 mg/kg for the first five postoperative days and 5 mg/kg per day over the follow up period of four weeks; mean plasma concentration: 421±64.7 μg/l; n=18) and methylprednisolone (2 mg/kg).

### Organ preparation and histology

Animals were sacrificed 4 weeks after transplantation. Hearts were processed for histology as recently described^8^. For the infarct characterization hearts from animals that had to be euthanized during the course of this study or previous studies^4,8^ were used. In brief, hearts were harvested and fixed in formalin for two days. Hearts were cut into 4 cross-sections and paraffin-embedded for histology. Dystrophin stained sections were used to determine infarct size and graft area. Human grafts were identified by human Ku80 staining. Masson Trichrome staining was used to discriminate the epicardial apposition from the scar tissue. Antigen retrieval and antibody dilution combinations were used as recently reported^8^.

### Statistical Analysis

Statistical analyses were performed with GraphPad Prism 9, USA. Comparison among two groups was made by two-tailed unpaired Student’s t-test. When two factors affected the result (e.g. time point and group), 2-way ANOVA analyses and Tukey’s-Test for multiple comparisons were performed. Error bars indicate SEM. P-values are displayed graphically as follows: * p < 0.05, ** p < 0.01, *** p < 0.001.

## Results

### EHT generation

The aim of this study was to evaluate whether EHT-patch transplantation in a chronically injured heart model demonstrates similar beneficial effects as at the subacute stage of injury^4,8^. EHT patch transplantation remuscularized the injured heart in a dose-dependent manner in a subacute model. However, only the highest dose (15×10^6^ cells per EHT patch) improved left ventricular function^8^. Accordingly, EHT patches containing 15×10^6^ cells were generated (troponin t positivity 76±2.6%; n=9) and used for the current study. EHT patches started to beat coherently after 5 days in culture and were cultured for 21 days prior to transplantation. At the time of implantation all EHTs demonstrated a coherent beating pattern.

### Remuscularization of the chronically injured heart

Myocardial injury was induced by a liquid nitrogen-cooled probe. Four weeks after injury induction, the cryo-lesion was covered with EHT patches (n=9) or cell-free patches (n=7). Hearts were harvested 28 days after transplantation for histological analysis (Figure 1A). Histomorphometric analysis demonstrated large transmural scars (Figure 1B) amounting to 25±1.7% of the left ventricle in the EHT patch group (n=9) and 28±3.7% (n=7; Figure 1C) in the cell-free patch group. Human grafts were identified by human Ku80 immunostaining (Figure 1B). Eight out of nine hearts in the EHT patch group contained engrafted human CMs. Graft size was 5±1.3% of the total scar area (Figure 1D). One heart was excluded from this analysis because it was not integrated in but only loosely connected to the heart, even though the EHT patch survived transplantation. Hearts that were transplanted with cell-free control patches showed epicardial fibrotic appositions in contrast compared to the intervention group (Supplemental Figure 1A). Cell counting of human Ku80^+^ cells in the graft amounted to 914±217 cells/section (Figure 1E; Supplemental Figure 1B).

**Figure 1.**
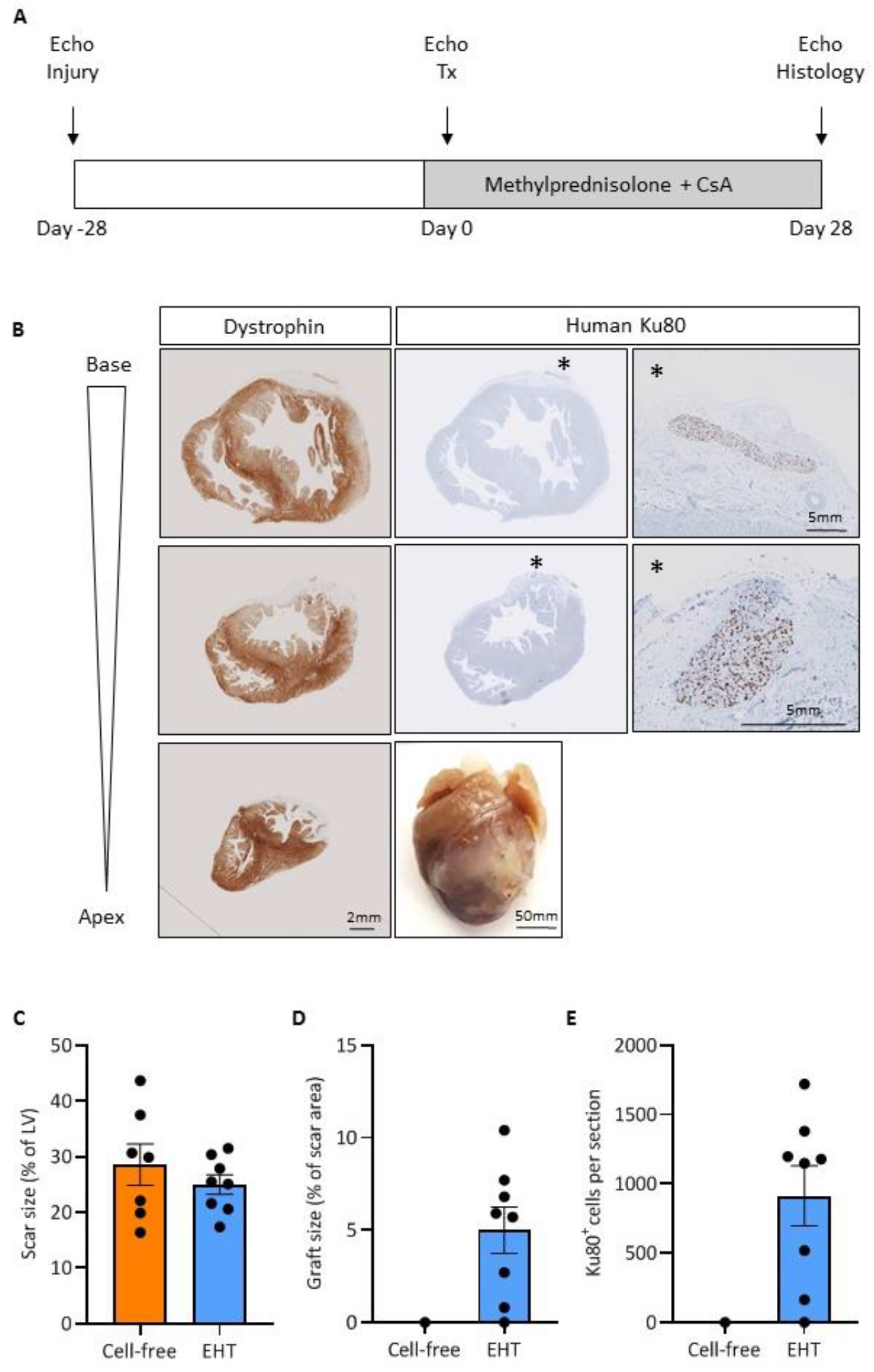
Histological evaluation of guinea pig hearts four weeks after transplantation. **A)** Experimental protocol. **B)** Picture of a heart four weeks after EHT patch transplantation and dystrophin- and human Ku80-stained sections of the respective heart, graft structures marked with an asterisk are shown at higher magnification. **C)** Quantification of scar size analysis and **D)** graft size. **E)** Quantification of Ku80^+^ cells per section. Mean ± SEM values are shown. CsA indicates Cyclosporine A; LV, left ventricle.

Grafts were surrounded by scar tissue without direct connection to viable host myocardium. Histological analysis exhibited that grafts consisted of densely aligned cardiomyocytes (cell density: 3233±112 cells per mm^2^; n=3) that mainly expressed the ventricular isoform of myosin light chain (MLC2v 98.6±0.8%; Figure 2A) with regular sarcomere structure. Sarcomere length of engrafted cardiomyocytes did not differ from host myocardium (1.9±0.0 μm vs. 1.9±0.0 μm; n=3, respectively; Figure 2B). Four weeks after transplantation grafts were vascularized by host-derived vessels (negative for human Ku80; Supplemental Figure 1C). Vascular density the graft was lower than in the remote myocardium (203±18.3 vessels per mm^2^ vs. 744±80.6 vessels per mm^2^ in the host myocardium; n=3; Figure 2C).

**Figure 2.**
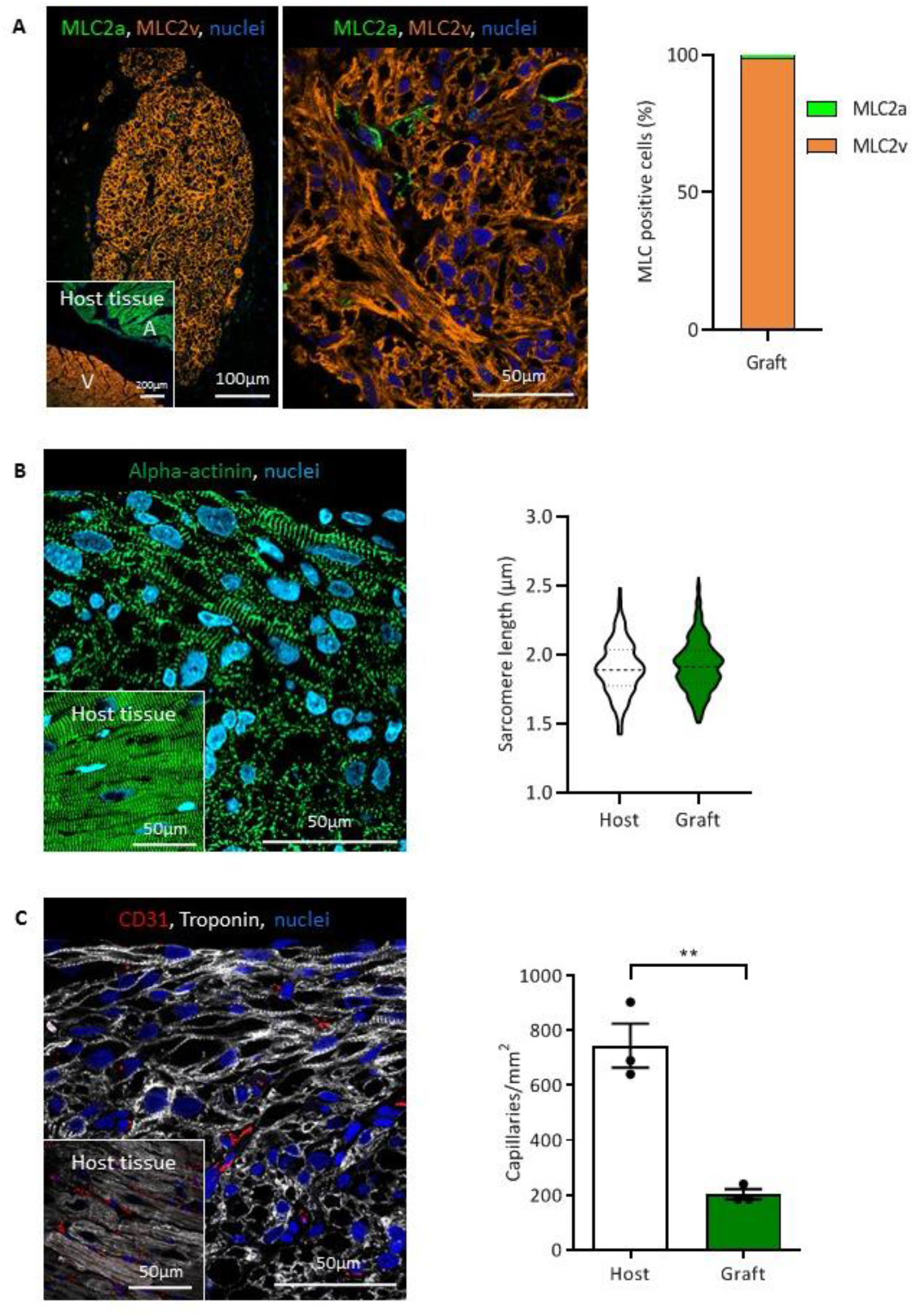
Histological analysis of engrafted cardiomyocytes. **A)** Myosin light chain staining of the atrial (MLC2a) and ventricular (MLC2v) isoform and quantification. **B)** Alpha-actinin staining and quantification of sarcomere length in human grafts compared to guinea pig host myocardium (n=3 hearts/>30 sarcomeres per graft or host myocardium). **C)** Human graft four weeks after transplantation and host myocardium stained for troponin and CD31. Quantification of graft vascularization four weeks after transplantation (n=3 hearts/3 high magnification images per graft or host myocardium). Each data point represents one heart. Mean ± SEM values are shown, **P<0.01. A indicates Atrium; V, Ventricle.

Cardiomyocyte proliferation after transplantation has been shown to participate in graft development^8^. Therefore, cardiomyocyte cell cycle activity was assessed by double-immunostaining for Ki67 and human Ku80. Four weeks after transplantation 10±2.2% engrafted human cells were still in the cell cycle (Figure 3A). To assess whether EHT patch transplantation could inhibit the chronic remodeling process of the remote myocardium host cardiomyocyte size was assessed by WGA (wheat germ agglutinin) staining. Four weeks after transplantation cardiomyocyte area was not different between animals that received EHT patches compared to animals in the cell-free control patch group (236±4.0 μm^2^ vs. 242±4.3 μm^2^; n=3; Figure 3B).

**Figure 3.**
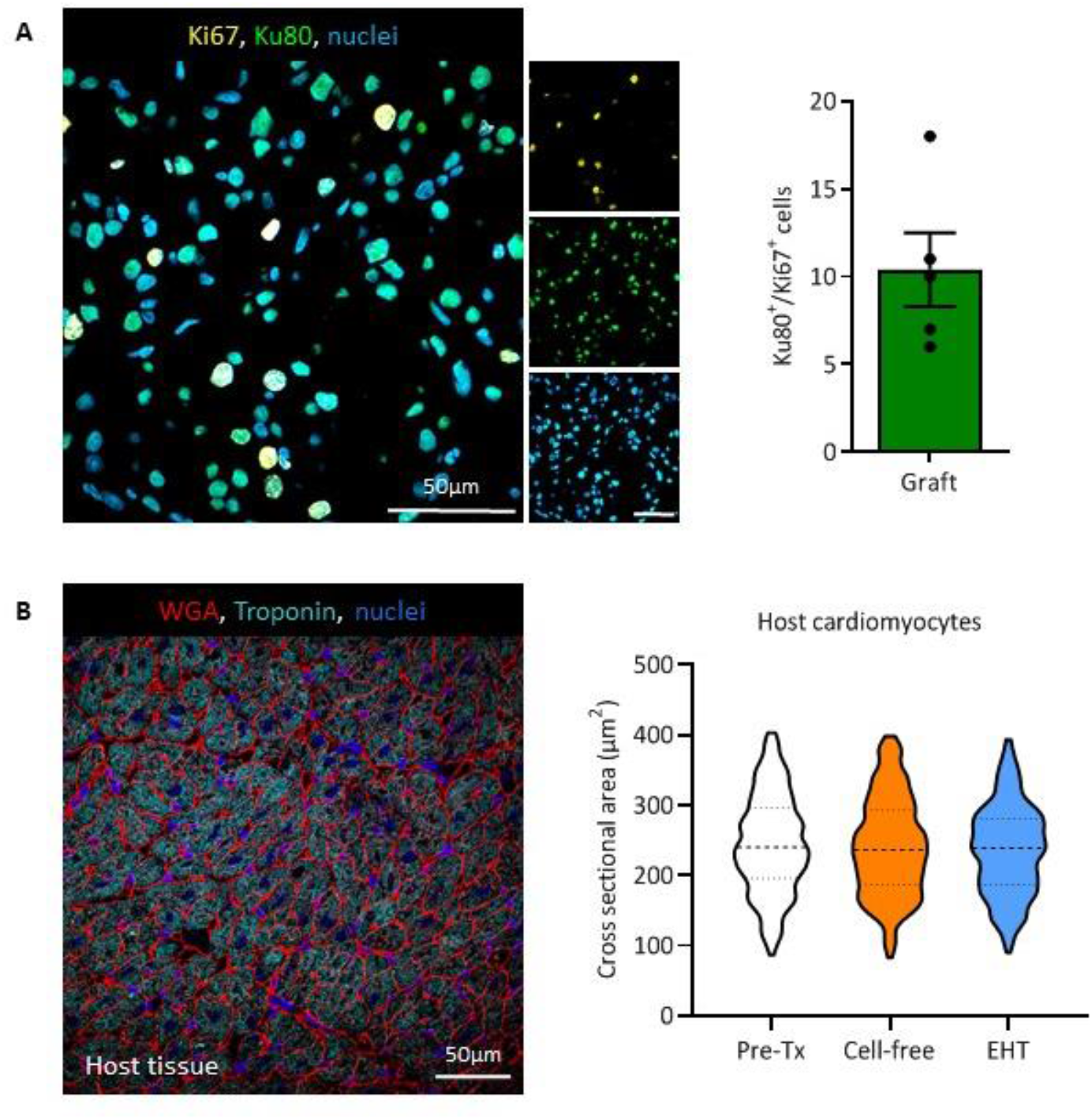
Analysis of graft proliferation and cell size in the remote myocardium. **A)** Human Ku80/Ki67 double immunostaining of a human graft four weeks after transplantation; Ki67/Ku80 positive nuclei (4 weeks after transplantation: n=2062 nuclei/N=5 hearts). **B)** Membrane (WGA) and troponin labeling of the remote myocardium and analysis of cell size before transplantation and four weeks after transplantation in the respective groups (n=3 hearts/3 high magnification images per remote myocardium). Mean ± SEM are shown. Pre-Tx indicates before transplantation.

### Functional consequences on left ventricular function

Functional consequence of EHT patch transplantation was evaluated by serial echocardiography (Figure 1A and 4A). Fractional area change (FAC) at baseline was 46±2.1%. Myocardial injury resulted in a significant reduction of left ventricular function four weeks after injury (FAC; control group: 31±3.6% vs. EHT group 28±3.5%). EHT patch transplantation did not significantly improve left ventricular function (FAC 36±4.4%) compared to the animals of the control group (FAC 34±3.3%; Figure 4B).

**Figure 4.**
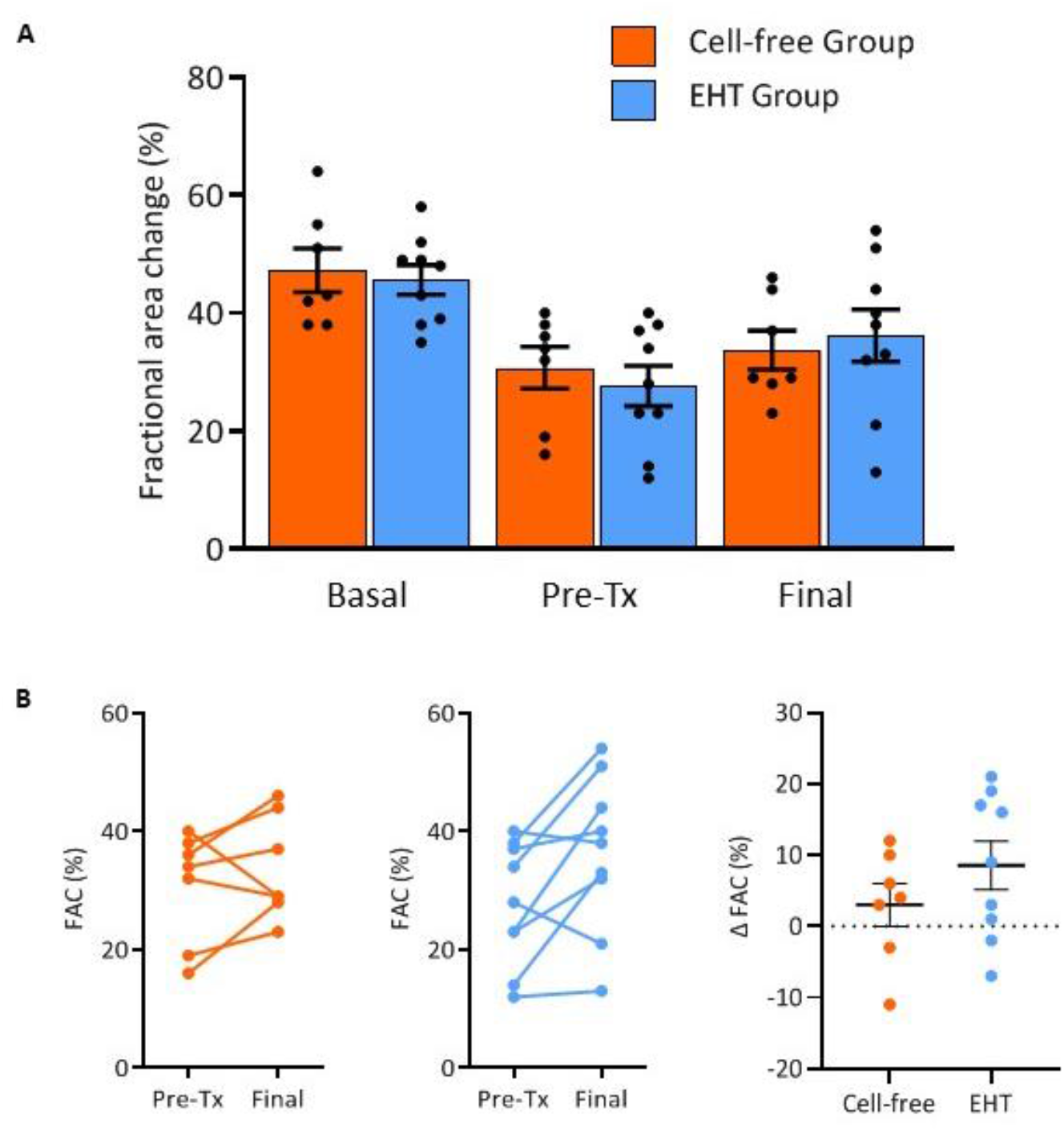
Echocardiographic evaluation. **A)** Fractional area change (FAC) analysis at baseline, four weeks post-injury and four weeks after transplantation (n=7 cell-free group, n=9 EHT patch group). **B)** Differences in FAC between day 28 (post-injury) and four weeks after transplantation. Mean ± SEM are shown. Pre-Tx indicates before transplantation; EHT, engineered heart tissue.

### Scar development over time

Given the striking difference between engraftment rates in the current study compared to the transplantation in a subacute setting^8^, we investigated scar development over time. Early after injury, the scars were clearly demarcated by inflammatory cells in the scar border zone (Figure 5). Seven days after injury, which represents the time point at which EHT patches were transplanted in a subacute setting, hematopoietic cells were visible throughout the scar, whereas they were almost absent 28 days after injury (Supplemental Figure 2A). In contrast, extracellular matrix (ECM) showed an opposite kinetic. Masson trichrome staining revealed first ECM deposition three days after injury. It increased over time and was paralleled by a thinning of the anterior ventricular wall (Figure 5) that showed a low vascularity (Supplemental Figure 2B).

**Figure 5.**
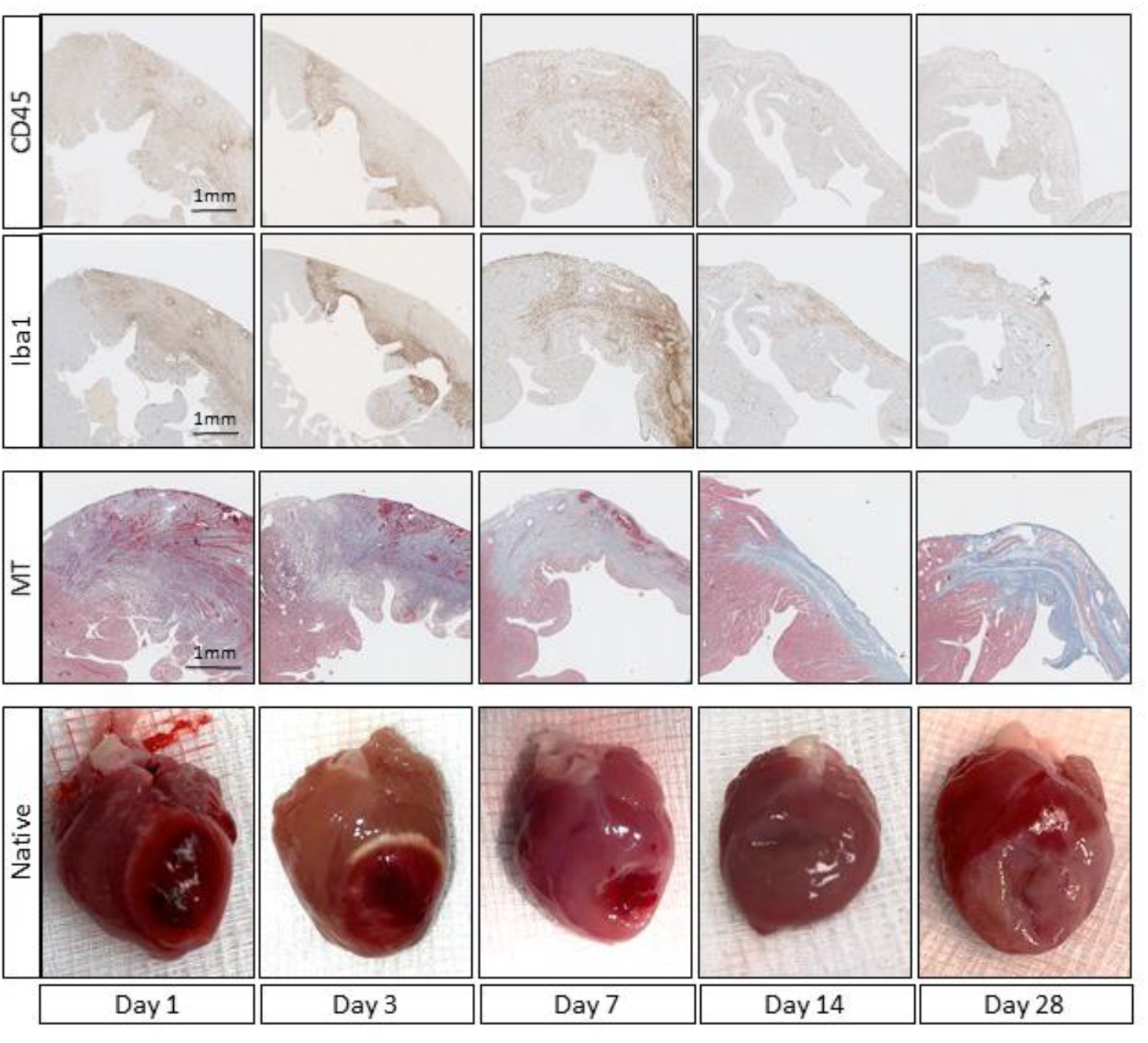
Characterization of myocardial infarction after cryo-injury. Immunohistochemical analysis of all hematopoietic cells (CD45) and macrophages (Iba1) at the site of injury and Masson Trichrome (MT) stained sections of the respective hearts.

## Discussion

In this study we evaluated EHT patch transplantation in a chronic injury guinea pig model. Recent studies in a subacute setting demonstrated that EHT patches remuscularized the injured heart in a dose-dependent manner and improved left ventricular function^8^. Similar approaches are currently evaluated in late preclinical and first clinical trials for patients with advanced heart failure (e.g. BioVAT-HF^12^). Yet, congestive heart failure most often represents a chronic remodeling process that, besides a loss of cardiomyocytes, involves cardiomyocyte hypertrophy and fibrosis. Evidence from neuroscience indicates chronically injured organs represent more difficult targets for regenerative strategies^13,14^. Similarly, cardiomyocyte injection, which has repeatedly demonstrated beneficial effects in subacute injury models, resulted in smaller grafts and demonstrated no beneficial effect on LV function in chronic injury models^10,15^. Yet, it was unknown whether EHT patch transplantation might represent a strategy to repair the chronically injured heart.

In the chronic injury model of this study, left ventricular function was lower at the time of transplantation (four weeks after injury) than in a previous subacute model^8^ (one week after injury), indicating a chronic adverse remodeling process. EHT patch transplantation resulted in a partial remuscularization of the chronically injured heart, but did not improve left ventricular function compared to animals that received cell-free patches. When comparing the results from the current study to the results from a transplantation study in a subacute setting^8^, two main differences are apparent: i) human grafts were smaller (5% vs. 12%) when transplanting the same hiPSC-CM EHTs and ii) EHT patch transplantation did not improve left ventricular function. Given that our previous study^8^ also did not observe significant improvement of left ventricular function at a cell/patch dose that reached 5% graft size, the data indicate that the smaller graft size per cell input in the chronic injury model simply did not suffice to exert a relevant effect on heart function. Similarly, a study in which cardiac constructs were transplanted four weeks after injury also reported no significant improvement in left-ventricular function^16^.

Scar size did not differ between the subacute^8^ and the chronic injury model, indicating that the lower degree of remuscularization was not due to an increase in injury area but to smaller engraftment after transplantation. This hypothesis is further substantiated when comparing the number of human Ku80^+^ nuclei per section (2250±680 subacute vs. 914±217 chronic), which leaves the question which factors cause the lower engraftment rate.

Factors that might influence engraftment include the following: i) Vascularization. Blood vessels might act as guiding structures for cardiomyocytes during regeneration^17^ and, maybe more importantly, deliver oxygen and nutrients to the transplanted cells. Four weeks after transplantation, graft vascularization was lower than in the host myocardium but did not differ between grafts in the subacute versus chronic model. However, we cannot rule out that vascularization during the first days after transplantation, i.e. the time when oxygen supply is critical was lower. Indeed, blood vessel density at the injury site was lower 4 weeks than 1 week after injury (Supplemental Figure 2B). In contrast to a study that evaluated direct intramyocardial cell injection in a chronic injury model^15^, we did not find evidence for human endothelial cells to participate in graft development. ii) Cardiomyocyte proliferation. Cardiomyocyte proliferation participates in graft formation^8,18,19^ and a smaller graft size could be simply explained by a lower number of proliferating cardiomyocytes. Yet, cell cycle activity did not differ between the grafts in a subacute and chronic model. Similarly to vascularization, differences in the proliferation dynamics at an early time point after transplantation might have gone unnoticed.

There are two more factors that most likely influence cell transplantation success: i) inflammation and ii) scar maturation. There is growing evidence that inflammation plays a pivotal role in cardiac regeneration. Species that show a profound regenerative response to cardiomyocyte injury, e.g. zebrafish, require immune cells to regenerate the heart and macrophage depletion minimizes their regenerative capacity^20^. Macrophage depletion also substantially blunted the regenerative response in neonatal mice^21^. Interestingly in the context of our study, the beneficial effects after bone marrow cell injection into the injured heart were recently shown to rely on an acute inflammatory response^22^. The cryoinjury intervention causes thermic damage to the guinea pig myocardium accompanied by sterile inflammation. We observed an influx of hematopoietic cells similar to murine LAD-ligation models^23^. Leukocyte invasion was seen as early as three days after injury and peaked around day 7. Four weeks after injury, the time point at which EHT patches were transplanted in the current study, the number of hematopoietic cells had already declined. Even though our study is not able to answer whether inflammation supports cell engraftment, it lays the foundation for future studies to directly address this question. Additionally, the lower engraftment rate could be explained by scar maturation. The initial injury is followed by a phase in which granulation tissue is gradually replaced by scar tissue over time. In a subacute setting seven days after injury, which reflects the time of transplantation in many studies, the site of injury has been invaded by fibroblasts that became activated and deposit ECM proteins. Yet, in the subacute setting, scars are not fully matured^24,25^. Over time collagen deposition increases^26^ and fibroblasts undergo morphological adaptations (e.g. alignment) and change in gene expression towards a tendon-like state occurs^25^. These maturation processes might also negatively impact cell engraftment.

In summary, this study showed that EHT patches survive transplantation in a chronically injured heart. Yet, it also demonstrated that engraftment rate was lower highlighting the difficulties towards a clinical application.

## Acknowledgements

We thank Kristin Hartmann (UKE, Mouse Pathology Core Facility) for technical assistance with immunohistochemistry. We would also like to thank the laboratory animal facility staff, UKE for their support during the study.

## Author contributions

C.v.B. performed experiments, analyzed data and prepared the manuscript. A.S. performed cell differentiation and generated EHT patches, B.G. performed experiments (echocardiography and surgeries) and analyzed data, E.Q., J.S., M.K., T.S. performed experiments, A.H., B.H. supervised the project and designed experiments, T.E., F.W. designed and supervised the project, acquired funding and prepared the manuscript.

## References

1. Khan, M. A. et al. Global Epidemiology of Ischemic Heart Disease: Results from the Global Burden of Disease Study. Cureus 12, (2020).

2. Hashimoto, H., Olson, E. N. & Bassel-Duby, R. Therapeutic approaches for cardiac regeneration and repair. Nature Reviews Cardiology 15, 585–600 (2018).

3. Söderlund, C. & Rådegran, G. Immunosuppressive therapies after heart transplantation — The balance between under- and over-immunosuppression. Transplant. Rev. 29, 181–189 (2015).

4. Weinberger, F. et al. Cardiac repair in Guinea pigs with human engineered heart tissue from induced pluripotent stem cells. Sci. Transl. Med. 8, 363ra148–363ra148 (2016).

5. Shiba, Y. et al. Human ES-cell-derived cardiomyocytes electrically couple and suppress arrhythmias in injured hearts. Nature 489, 322–325 (2012).

6. Laflamme, M. A. et al. Cardiomyocytes derived from human embryonic stem cells in pro-survival factors enhance function of infarcted rat hearts. Nat. Biotechnol. 25, 1015–1024 (2007).

7. Liu, Y. W. et al. Human embryonic stem cell-derived cardiomyocytes restore function in infarcted hearts of non-human primates. Nat. Biotechnol. 36, 597–605 (2018).

8. Querdel, E. et al. Human Engineered Heart Tissue Patches Remuscularize the Injured Heart in a Dose-Dependent Manner. Circulation (2021). doi:10.1161/CIRCULATIONAHA.120.047904

9. Pecha, S. et al. Human iPS cell-derived engineered heart tissue does not affect ventricular arrhythmias in a guinea pig cryo-injury model. Sci. Rep. 9, 1–12 (2019).

10. Shiba, Y. et al. Electrical Integration of Human Embryonic Stem Cell-Derived Cardiomyocytes in a Guinea Pig Chronic Infarct Model. J. Cardiovasc. Pharmacol. Ther. 19, 368–381 (2014).

11. Breckwoldt, K. et al. Differentiation of cardiomyocytes and generation of human engineered heart tissue. Nat. Protoc. 12, 1177–1197 (2017).

12. Stefan Weller. Start of First Clinical Trial on Tissue Engineered Heart Repair. idw (2021). Available at: https://idw-online.de/de/news762811. (Accessed: 17th June 2021)

13. Nagoshi, N., Okano, H. & Nakamura, M. Regenerative therapy for spinal cord injury using iPSC technology. Inflammation and Regeneration 40, (2020).

14. Stevens, K. R. & Murry, C. E. Cell Stem Cell Forum Human Pluripotent Stem Cell-Derived Engineered Tissues: Clinical Considerations. doi:10.1016/j.stem.2018.01.015

15. Fernandes, S. et al. Human embryonic stem cell-derived cardiomyocytes engraft but do not alter cardiac remodeling after chronic infarction in rats. J. Mol. Cell. Cardiol. 49, 941–949 (2010).

16. Riegler, J. et al. Human engineered heart muscles engraft and survive long term in a rodent myocardial infarction model. Circ. Res. 117, 720–730 (2015).

17. Marín-Juez, R. et al. Fast revascularization of the injured area is essential to support zebrafish heart regeneration. Proc. Natl. Acad. Sci. U. S. A. 113, 11237–11242 (2016).

18. El-Nachef, D. et al. Engrafted Human Induced Pluripotent Stem Cell-Derived Cardiomyocytes Undergo Clonal Expansion in Vivo. Circulation 143, 1635–1638 (2021).

19. Zhang, Y. et al. Dedifferentiation and proliferation of mammalian cardiomyocytes. PLoS One 5, 1–13 (2010).

20. Lai, S. L. et al. Reciprocal analyses in zebrafish and medaka reveal that harnessing the immune response promotes cardiac regeneration. Elife 6, (2017).

21. Lavine, K. J. et al. Distinct macrophage lineages contribute to disparate patterns of cardiac recovery and remodeling in the neonatal and adult heart. Proc. Natl. Acad. Sci. U. S. A. 111, 16029–16034 (2014).

22. Vagnozzi, R. J. et al. An acute immune response underlies the benefit of cardiac stem cell therapy. Nature 577, 405–409 (2020).

23. Virag, J. I. & Murry, C. E. Myofibroblast and Endothelial Cell Proliferation during Murine Myocardial Infarct Repair. Am. J. Pathol. 163, 2433–2440 (2003).

24. Yokota, T. et al. Type V Collagen in Scar Tissue Regulates the Size of Scar after Heart Injury. Cell 182, 545–562.e23 (2020).

25. Fu, X. et al. Specialized fibroblast differentiated states underlie scar formation in the infarcted mouse heart. J. Clin. Invest. 128, 2127–2143 (2018).

26. Rusu, M. et al. Biomechanical assessment of remote and postinfarction scar remodeling following myocardial infarction. Sci. Rep. 9, 1–13 (2019).

