## Supplemental Figures for "Human engineered heart tissue transplantation in a guinea pig chronic injury model"

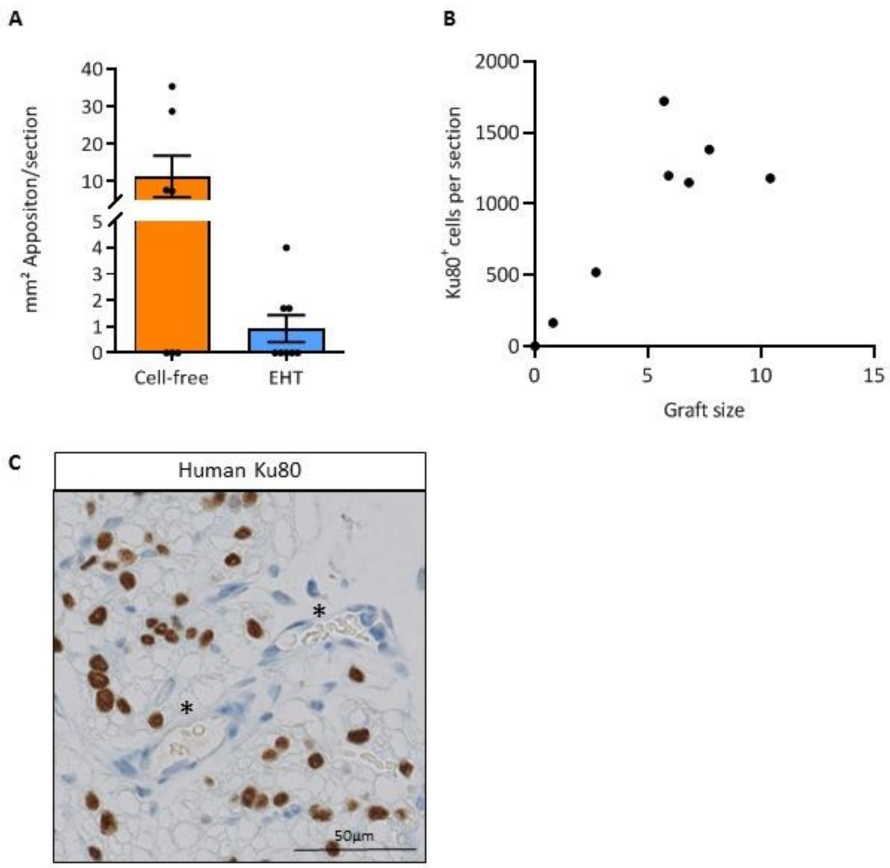


**Supplemental Figure 1. A)** Area based analysis of the epicardial apposition of fibrotic tissue in the respective groups (n=7 cell-free group, n=8 EHT patch group). **B)** Correlation between graft size four weeks after transplantation and Ku80 positive cells. **C)** Human graft stained for human Ku80. Asterisks mark host-derived (Ku80 negative) vessels. Mean ± SEM are shown. EHT indicates engineered heart tissue.

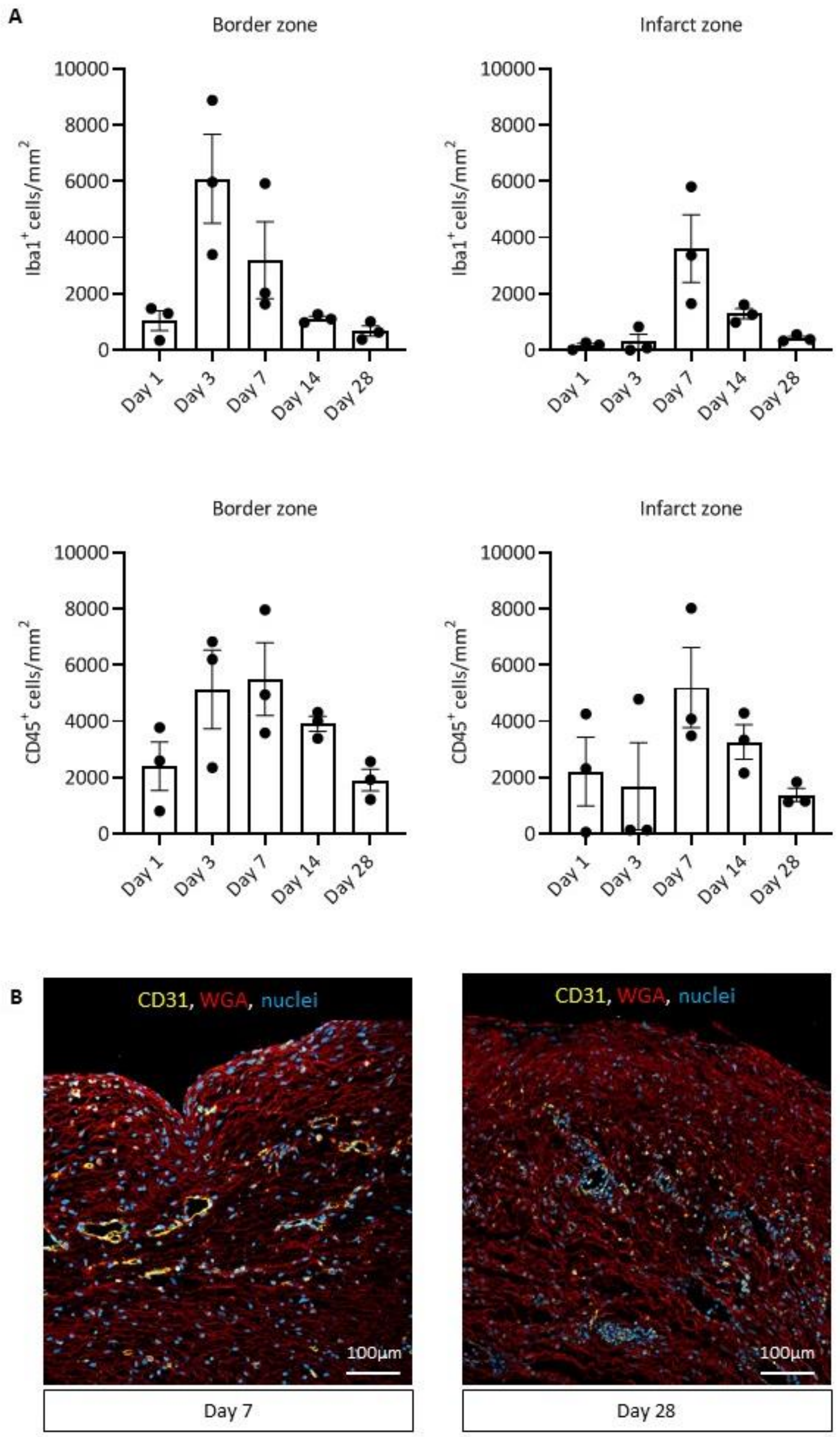


**Supplemental Figure 2. A)** Quantification of CD45$^+$ cells and macrophages (Iba1) at different time points after injury (n=3 hearts/3 high magnification images per heart). Each data point represents one heart. **B**) Membrane (WGA) and CD31 labeling of the infarct zone one week and four weeks after cryo-injury. Mean ± SEM values are shown.
